# Intelli-NGS: Intelligent NGS, a deep neural network-based artificial intelligence to delineate good and bad variant calls from IonTorrent sequencer data

**DOI:** 10.1101/2019.12.17.879403

**Authors:** Aditya Singh, Prateek Bhatia

## Abstract

**Background:** IonTorrent is a second-generation sequencing platform with smaller capital costs than Illumina but is also prone to higher machine error than later. Given its lower costs, the platform is generally preferred in developing countries where next-generation sequencing is still a very exclusive technique. There are many software tools available for other platforms but IonTorrent. This makes the already tricky analysis part more error-prone.

**Motivation:** We have been using the IonTorrent platform in our hospital setting for aiding diagnosis or treatment for the past couple of years. Given to our experience, analysis part of IonTorrent data takes the longest time and still, we used to get stuck with certain variants which seemed fine on looking at their metrics but were found to be negative in Sanger sequencing verification. This made us determined to develop a tool that could aid us in reducing false positive and negative rates while still retaining good recall. The artificial intelligence-based technique was our final choice after developing pipelines with less success.

**Methodology:** The artificial intelligence was developed from scratch in Python 3 using TensorFlow fully connected dense layers. The model takes VCF files as input and solves each variant based on the thirty-five parameters given by the IonTorrent platform, including the flow-space information which is missed by variant callers other than the default torrent variant caller.

**Results:** The final trained model was able to achieve an accuracy of 93.08% and a ROC-AUC of 0.95 with GIAB validation data. The additional program that was written to run the model annotates each variant using online databases such as dbSNP, ClinVar and others. A probability score for each outcome for each variant is also provided to aid in decision making.

**Availability:** The model and running code are available for free only for non-commercial users at https://www.github.com/aditya-88/intelli-ngs.

## 1 Introduction

Next-generation sequencing (NGS) has gradually become an indispensable tool for disease profiling, research and novel genetic markers discovery among other applications of the technology. Although, the costs of sequencing have gone down reasonably still these techniques are out of reach or exclusive to special cases in developing countries. IonTorrent next-generation sequencing platform has higher error rates than other platforms like Illumina^1^ but the former has certain advantages like lower capital costs and flexibility in targeted sequencing. We have been using IonTorrent GeneStudio S5 for the last couple of years and have run hundreds of patient samples and have observed the error rates of IonTorrent being on the higher side. When we are dealing with patient data, accuracy and precision of the results become of utmost importance. In the light of the same, we tried various other pieces of software among which, we found Genome Analysis Toolkit (GATK)^2^ with custom hard filters to improve the variant calls. But, this too, was either increasing the false-negative or false-positive rates. Given to the fact that any tool that we use other than the default variant caller, we will lose a very crucial piece of data termed as *“flow space data”* which incorporates the raw reads of change in voltage while the sequencer was running. This vital information helps in delineating good and bad variant calls and is available in the variant call format (VCF) file per variant outputted by the instrument. We hypothesized that, by using this information in conjunction with other metrics in the VCF file, we can train a machine learning model based on fully connected dense layers to delineate good and bad variant calls. A total of thirty-five parameters (supplementary table S1) were used as input layer for the machine learning model and two output layers each for good and bad variant calls with three hidden fully connected and three dropout layers in TensorFlow 2.0 using Keras backend. The model was developed in Python 3 along-with a wrapper code to use the model for analysing actual VCF files.

## 2 Materials and Methods

### 2.1 Preparation of samples for Machine learning

Standard IonTorrent sequenced BAM alignment file for sample accession number NA12878 from Genome in a Bottle (GIAB: https://github.com/genome-in-a-bottle/giab_data_indexes) consortia database was downloaded and uploaded to local IonReporter server running UI version 5.10.38 and set for variant calling using all default parameters. A short Python 3 code was used to extract the 35-variant metrics from the resulting file and a spreadsheet of the same was made in Microsoft Excel. A total of 1,336 confirmed negative calls listed on the website was selected out from the total variants as a negative subset and given code of number zero. A total of 36,198 variants falling in the high confidence regions which were confirmed to be good calls from multiple sequencing platforms by the consortia were selected as confirmed positive calls and given the code of number one. Since we ended up with around 27 times more positive variants than negatives, we needed to select the same number of positive variants that is, 1,336. In order to do so, we firstly removed any variant with less than 30 read depth as it is a cut off for germline variants, hence ending up with 33,757 positive variants. The final subset of 1,336 positive variants from the whole set of 33,757 variants was selected at random using Python module Pandas. The resulting file was exported to another Excel Sheet and used for training of the model. The training program takes both files and merges them together. After picking up the file for training, the model training program also codes the “type” field of the variant for SNP, MNP, INS, DEL, Complex as 0 to 4 providing the model with an additional weight that governs the type of variant. The final resulting variants were 2,672, out of which randomly 5% data was selected for validation per epoch and rest for training.

### 2.2 Development of the model

Python 3 along with TensorFlow, Pandas, Sci-kit learn, Numpy and Matplotlib modules were used to code the model. The model is based on artificial neural network (ANN) architecture which comprises of fully connected dense layers. The architecture of the model can be referred to in table 1 and figure 1 (created using https://github.com/lutzroeder/netron). The model was compiled with Adamax optimizer, sparse categorical cross-entropy as loss function with default learning rate.

**Table 1.** The architecture of the machine learning model

| S.No. | Type of layer | Units | Activation |
| --- | --- | --- | --- |
| 1. | Dense (input: 35,) | 35 | Sigmoid |
| 2. | Dropout | 0.4 | - |
| 3. | Dense | 150 | Sigmoid |
| 4. | Dropout | 0.4 | - |
| 5. | Dense | 35 | Sigmoid |
| 6. | Dropout | 0.4 | - |
| 7. | Dense | 35 | Sigmoid |
| 8. | Dropout | 0.4 | - |
| 9. | Dense (output) | 2 | SoftMax |

**Figure 1:**
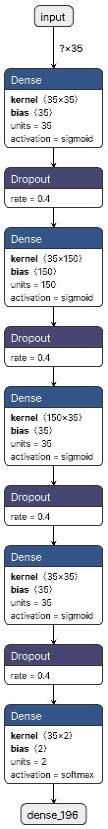
Architecture of the machine learning model

### 2.3 Development of the wrapper code

The wrapper code written to analyse actual VCF files using the developed model was also written in Python 3 with sub-modules Sys, OS, TensorFlow, Pandas, Sci-kit Allel and MyVariant. Additionally, bcftools^3^ was used to segregate multiple variants at one site during the pre-processing stage of the VCF files. The wrapper code is capable of accepting practically any number of VCF files at once and processes them one by one. The code creates a Microsoft Excel sheet pertaining to each VCF file with HGVS codes for all variants, prediction verdict of the machine learning model along with probabilities of both outcomes and annotation using available online databases enabled by Python MyVariant module.

### 2.4 Evaluation of the model

A Python 3 code was written to evaluate the efficacy of the model using GIAB data. The evaluator code was additionally comprised of TensorFlow, Pandas, Matplotlib, Seaborn, Scikitplot, Numpy and Sci-kit learn. An equal number of positive and negative variant calls from the whole data set was randomly selected and fed to the model for evaluation. Confusion matrix, cumulative gain, Kolmogorov-Smirnov (KS) test, Matthews correlation, lift curve and receiver operating curve (ROC) was calculated for the model evaluation.

## 3 Results

### 3.1 Model training

In the test dataset, the model reached the highest accuracy of 94.02% after 700 epochs with loss functions of 0.2258 and 0.1816 in training and validation dataset respectively. The loss function for validation data is less than that of the training data which signifies that the model is not hogged by overfitting. For the test set of 134 values, the model was able to have a recall of 100% with 61 true positives, 65 true negatives, 8 false positives and no false negatives.

### 3.2 Model evaluation

The model was evaluated with a total of 2,672 samples formed of an equal number of good and bad calls from the GIAB dataset chosen at random. In the evaluation set, the model was able to achieve a recall of 99.55% with 1,330 true positives, 1,157 true negatives, 179 false positives and 6 false negatives. In figure 2 and table 2, we can find various performance graphs and details for the model. From the ROC plot, we can infer that the micro average ROC area under the curve was 0.97 and individually for both classes, it was 0.95. The model performed well with a true positive rate of 0.9955, the negative predictive value of 0.9948, specificity of 0.866, precision of 0.8814 and overall accuracy of 93% with an F1 score of 0.935. The Matthews correlation coefficient for the model was 0.8688. The KS statistic test suggests the maximum segregation to be 86.4%, the cumulative gain graph shows a decent separation from the baseline and the lift curve suggests that the usage of the model greatly improves the predictability when compared with the random selection method.

**Table 2:** Evaluation report of the model

| Measure | Value | Derivations |
| --- | --- | --- |
| Sensitivity | 0.9955 | $TPR = TP / (TP + FN)$ |
| Specificity | 0.8660 | $SPC = TN / (FP + TN)$ |
| Precision | 0.8814 | $PPV = TP / (TP + FP)$ |
| Negative Predictive Value | 0.9948 | $NPV = TN / (TN + FN)$ |
| False Positive Rate | 0.1340 | $FPR = FP / (FP + TN)$ |
| False Discovery Rate | 0.1186 | $FDR = FP / (FP + TP)$ |
| False Negative Rate | 0.0045 | $FNR = FN / (FN + TP)$ |
| Accuracy | 0.9308 | $ACC = (TP + TN) / (P + N)$ |
| F1 Score | 0.9350 | $F1 = 2TP / (2TP + FP + FN)$ |
| Matthews Correlation Coefficient | 0.8688 | $TP*TN - FP*FN / \sqrt{(TP+FP)*(TP+FN)*(TN+FP)*(TN+FN)}$ |

• TPR: True positive rate; TP: True Positives; FN: False Negatives; SPC: Specificity; TN: True Negatives; FP: False Positives; PPV: Precision; NPV: Negative Predictive Value; FPR: False Positive Rate; FDR: False Discovery Rate; ACC: Accuracy

**Figure 2:**
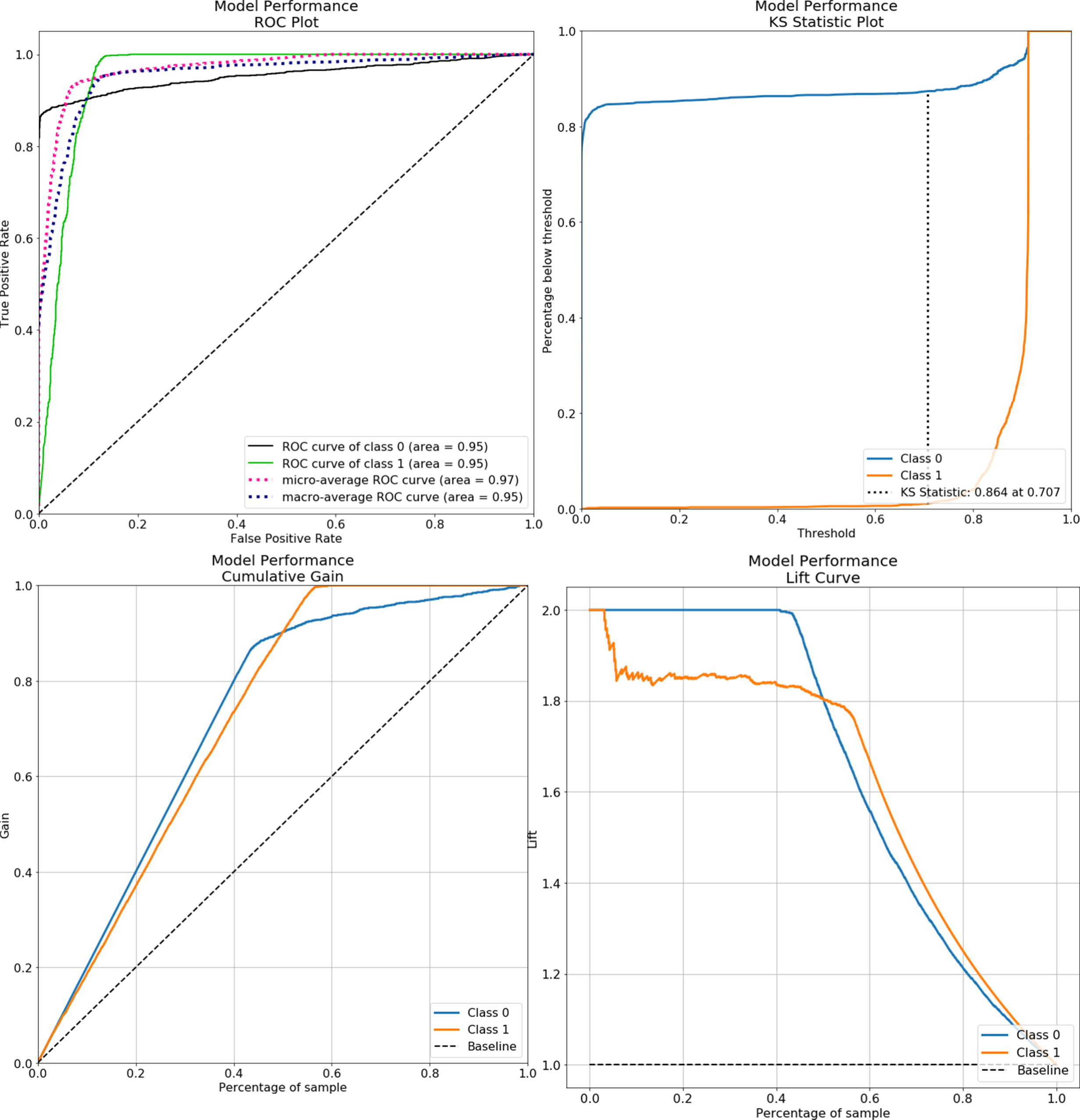
Model performance evaluation charts

## Discussion

The developed model gives out probability score and decisions for each variant call which augments the final decision-making process of accepting and rejecting variants. The model was developed in TensorFlow, which is an open-source and popular machine learning module and hence more users can fine-tune this model based on their own data using the transfer learning method of TensorFlow. The model with its wrapper code is merely 219 kilobases in size and can easily run on any modern-day POSIX compliant computer and a graphics card isn’t necessary for the same. A similar tool, GARFIELD-NGS was released in 2018 by Ravasio *et al.*,^4^ but the has much lower accuracy of 0.804 for INDELs and comparable accuracy of 0.955 for SNVs. Also, it utilizes different models for INDELs and SNVs, is incapable of multiplexing and is developed in H_2_O.ai, which is a commercial software. Intelli-NGS was developed with an additional field which scored different types of variations; hence the model can solve the problem by populating weights for the additional parameter. When we compare the architecture of the two models, Intelli-NGS solves the problem with a total of 12,862,500 connections while GARFIELD-NGS model for SNVs requires 972,000,000 and INDELs requires1,620,000,000 connections which is 75.57 and 125.95 folds respectively more complex than the former and hence will consume more computing resources to run. Apart from this, there are a few other tools like DeepVariant^5^, SNPSVM^6^ and GotCloud^7^ which combine variant calling and filtration in one algorithm, this generally requires workstation level computer and changes the whole pipeline of work, unlike the tools which work with the already produced VCF files. The lightweight model of Intelli-NGS can effectively delineate good and bad variants from the IonTorrent data along with probability scores, aiding in the decision-making process while scrutinizing variants. Also, the accompanying code written to utilize the model is capable of multiplexing samples and loads the model into the memory only once per session, unlike the other discussed code which requires unique sessions for each sample and hence will load and unload the model each time to and from the memory, ultimately consuming more resources and time. The code additionally also annotates each variant with available databases online using MyVariant module of Python 3. All the code written and utilized are open source and hence promotes co-development and better understanding. The code requires no compilation or advanced installation of tools, almost all dependencies are available from Python’s repository but bcftools, which is generally available in all major Linux repositories and in the Homebrew (https://brew.sh) repository of macOS.

## Conclusion

Intelli-NGS was developed utilizing the high confidence GIAB data and tuned to use the lowest amount of complexity in architecture enabling faster operations on even average computers. The model has an overall accuracy of 93.08%, recall of 99.55%, a specificity of 86.60%, precision of 88.14% and negative prediction of 99.48%.

## Author Contributions

AS analysed the data, developed the artificial intelligence and evaluated the model. AS wrote the article. PB proofread the article with significant inputs.

