## supplementary table S1 for "Intelli-NGS: Intelligent NGS, a deep neural network-based artificial intelligence to delineate good and bad variant calls from IonTorrent sequencer data"

**Supplementary Table S1:** List of parameters used for training the machine

| **S.No.** | **Parameter** | **Description** |
| --- | --- | --- |
| 1. | AF | Allele frequency based on Flow Evaluator observation counts |
| 2. | AO | Alternate allele observations |
| 3. | DP | Total read depth at the locus |
| 4. | FAO | Flow Evaluator Alternate allele observations |
| 5. | FDP | Flow Evaluator read depth at the locus |
| 6. | FDVR | Level of Flow Disruption of the alternative allele versus reference. |
| 7. | FRO | Flow Evaluator Reference allele observations |
| 8. | FSAF | Flow Evaluator Alternate allele observations on the forward strand |
| 9. | FSAR | Flow Evaluator Alternate allele observations on the reverse strand |
| 10. | FSRF | Flow Evaluator Reference observations on the forward strand |
| 11. | FSRR | Flow Evaluator Reference observations on the reverse strand |
| 12. | FWDB | Forward strand bias in prediction. |
| 13. | FXX | Flow Evaluator failed read ratio |
| 14. | HRUN | Run length: the number of consecutive repeats of the alternate allele in the reference genome |
| 15. | LEN | Allele length |
| 16. | MLLD | Mean log-likelihood delta per read. |
| 17. | PB | Bias of relative variant position in reference reads versus variant reads. Equals Mann-Whitney U rho statistic P(Y>X)+0.5P(Y=X) |
| 18. | PBP | P-val of relative variant position in reference reads versus variant reads. Related to GATK ReadPosRankSumTest |
| 19. | QD | QualityByDepth as 4*QUAL/FDP (analogous to GATK) |
| 20. | QUAL | Quality score of the alignment |
| 21. | RBI | Distance of bias parameters from zero. |
| 22. | REFB | Reference Hypothesis bias in prediction. |
| 23. | REVB | Reverse strand bias in prediction. |
| 24. | RO | Reference allele observations |
| 25. | SAF | Alternate allele observations on the forward strand |
| 26. | SAR | Alternate allele observations on the reverse strand |
| 27. | SRF | Number of reference observations on the forward strand |
| 28. | SRR | Number of reference observations on the reverse strand |
| 29. | SSEN | Strand-specific-error prediction on negative strand. |
| 30. | SSEP | Strand-specific-error prediction on positive strand. |
| 31. | SSSB | Strand-specific strand bias for allele. |
| 32. | STB | Strand bias in variant relative to reference. |
| 33. | STBP | Pval of Strand bias in variant relative to reference. |
| 34. | TYPE | The type of allele, either snp, mnp, ins, del, or complex. Coded as 0 to 4 |
| 35. | VARB | Variant Hypothesis bias in prediction. |
